# MRI-assessed locus coeruleus contrast and functional response are not associated in young and late middle-aged individuals

**DOI:** 10.1101/2023.01.16.524213

**Authors:** Alexandre Berger, Ekaterina Koshmanova, Elise Beckers, Roya Sharifpour, Ilenia Paparella, Islay Campbell, Nasrin Mortazavi, Fermin Balda, Yeo-Jin Yi, Laurent Lamalle, Laurence Dricot, Christophe Phillips, Heidi IL Jacobs, Puneet Talwar, Riëm El Tahry, Siya Sherif, Gilles Vandewalle

## Abstract

The brainstem locus coeruleus (LC) influences a broad range of brain processes, including cognition. The so-called LC contrast is an accepted marker of the integrity of the LC that consists of a local hyperintensity on specific Magnetic Resonance Imaging (MRI) structural images. The small size of the LC has, however, rendered its functional characterization difficult in humans, including in aging. A full characterization of the structural and functional characteristics of the LC in healthy young and late middle-aged individuals is needed to determine to potential roles of the LC in different medical conditions. Here, we wanted to determine whether the activation of the LC in a mismatch negativity task changes in aging and whether the LC functional response was associated to the LC contrast. We used Ultra-High Field (UHF) 7-Tesla functional MRI (fMRI) to record brain response during an auditory oddball task in 53 healthy volunteers, including 34 younger (age: 22.15y ± 3.27; 29 women) and 19 late middle-aged (age: 61.05y ± 5.3; 14 women) individuals. Whole-brain analyses confirmed brain responses in the typical cortical and subcortical regions previously associated with mismatch negativity. When focusing on the brainstem, we found a significant response in the rostral part of the LC probability mask generated based on individual LC images. Although bilateral, the activation was more extensive in the left LC. Individual LC activity was not significantly different between young and late middle-aged individuals. Critically, while the LC contrast was higher in older individuals, the functional response of the LC was not associated with its contrast. These findings show that the age-related alterations of the LC structural integrity may not necessarily be related to changes in its functional response. The results further indicate that LC responses could remain stable in healthy individuals aged 20 to 70.

## INTRODUCTION

The Locus Coeruleus (LC) is a brainstem nucleus in the reticular formation that is gaining a lot of scientific attention. Being the main source of norepinephrine (NE) in the brain, this nucleus sends monosynaptic projections to almost all brain regions (Aston-Jones et al., 1986; Keren et al., 2009), forming the LC-NE system. The LC is involved in numerous processes including the regulation of anxiety and alertness, the gating of sleep, the detection of salient changes in the environment, and cognition over various domains (Sara, 2009). Alteration in the LC has been suggested in an array of psychiatric and neurological diseases including Alzheimer’s Disease (AD) dementia (Jacobs et al., 2021), Parkinson’s disease (Braak et al., 2003), rapid eye movement sleep behavior disorder (García-Lorenzo et al., 2013), insomnia disorders (van Someren et al., 2020), pathological anxiety (Morris et al., 2020), late-life major depression (Guinea-Izquierdo et al., 2021) or schizophrenia (Mäki-Marttunen et al., 2020).

*In vivo* investigations of the functions and structure of the LC have been hampered due to the small size of the nucleus, which consists of a pair of ∼15 mm long cylinder-shaped nuclei of 2.5 mm diameter with a total of ∼50.000 neurons in humans (Aston-Jones et al., 1986; Keren et al., 2009). Part of this difficulty was overcome using structural Magnetic Resonance Imaging (MRI) sequences seemingly sensitive to the high content of neuromelanin within the LC. Using such sequences, the LC appears hyperintense and can be isolated from its surrounding tissues (Kelberman et al., 2020). The recent development of Ultra-High Field (UHF) 7-Tesla (7T) MRI further permitted submillimeter higher resolution and higher signal-to-noise ratio for imaging the LC (Priovoulos et al., 2018). The biophysical origin of the LC contrast and the significance of its variations remain debated (Galgani et al., 2021; Priovoulos et al., 2020). The consensus is, however, that it is related to the structural integrity of the cells constituting the LC. Studies assessing LC functional responses remain scarce, likely because of the difficulty to reliably detect LC activation using functional MRI (fMRI) in humans.

In the context of healthy aging, studies reported that the LC contrast increases in adulthood up to about 60 years to decline thereafter (Jacobs et al., 2021; Liu et al., 2019; Shibata et al., 2006). The variability of the LC contrast in aging may be related to the presence of AD pathology, altering the structural integrity of the LC (Jacobs et al., 2021). A recent *in vivo* 3T fMRI study that used a visual novelty paradigm in humans further reported that lower activity and functional connectivity of the LC were associated with amyloid-related cognitive decline in cognitively unimpaired older individuals (Prokopiou et al., 2022a). However, whether LC functional response changes with healthy aging is not established. Likewise, whether age-related changes in LC contrast is associated with a detectable change in its functional response has not been investigated.

Here, we first tested whether the functional response of the LC during an auditory oddball task – a robust paradigm used to recruit the LC (Murphy et al., 2014) – differed between young and late middle-aged healthy individuals. Based on previous electrophysiology and fMRI studies that reported a reduced brain response to oddball events in aging (Juckel et al., 2012; van Dinteren et al., 2014), we expect a more prominent response in the LC of younger adults. We further investigated whether the LC response was related to the LC integrity, as assessed through its contrast, to determine if a structural-functional relationship may exist in the nucleus. Overall, this work aims to further our current knowledge about the effect of healthy aging on LC physiology and anatomy. A better functional and structural characterization of this nucleus in healthy individuals could help to detect abnormal features in different medical conditions. This could help to improve the detection of LC-related disorders and diseases in young and late middle-aged individuals, better characterize their evolution and potentially help to develop therapies that are targeting the LC-NE system.

## MATERIALS AND METHODS

### Participants

A sample of 53 healthy participants of both sexes, composed of 34 healthy young (age: 22.15 ± 3.27 y, 29 women) and 19 late middle-aged (age: 61.05 ± 5.3 y, 14 women) individuals were included in this study. A summary of the demographic data can be found in **Table 1**. This study was approved by the faculty-hospital Ethics Committee of the University of Liège. All participants provided their written informed consent and received a financial compensation.

**Table 1:**
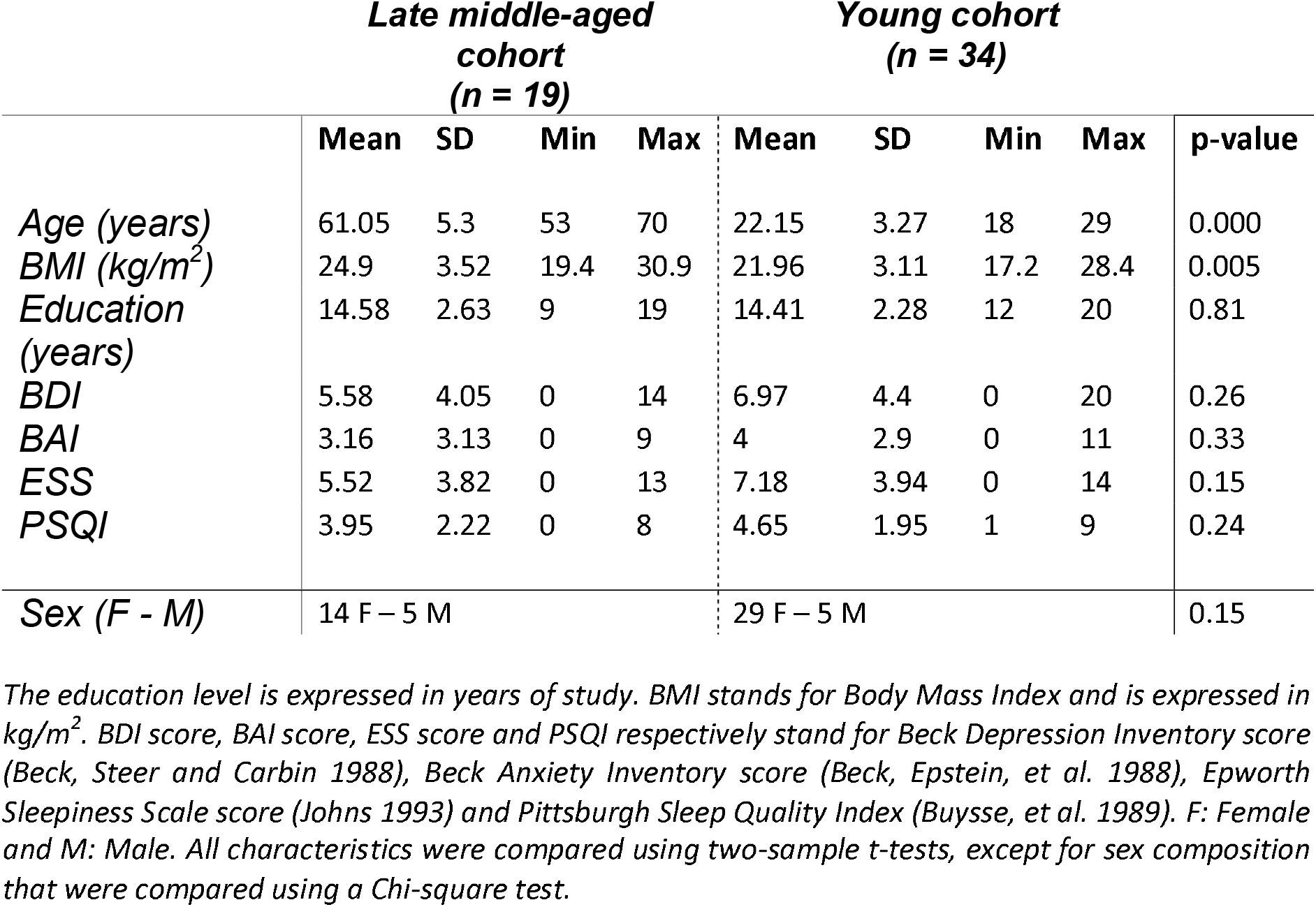
Characteristics of the **study samples**. The p-values shown in the table correspond to two- sample t-tests comparing the characteristics between the young and late middle-aged cohort.

|  | **Late middle-aged cohort**<br>**(n = 19)** |  |  |  | **Young cohort**<br>**(n = 34)** |  |  |  |  |
| --- | --- | --- | --- | --- | --- | --- | --- | --- | --- |
|  | Mean | SD | Min | Max | Mean | SD | Min | Max | p-value |
| *Age (years)* | 61.05 | 5.3 | 53 | 70 | 22.15 | 3.27 | 18 | 29 | 0.000 |
| *BMI (kg/m<sup>2</sup>)* | 24.9 | 3.52 | 19.4 | 30.9 | 21.96 | 3.11 | 17.2 | 28.4 | 0.005 |
| *Education (years)* | 14.58 | 2.63 | 9 | 19 | 14.41 | 2.28 | 12 | 20 | 0.81 |
| *BDI* | 5.58 | 4.05 | 0 | 14 | 6.97 | 4.4 | 0 | 20 | 0.26 |
| *BAI* | 3.16 | 3.13 | 0 | 9 | 4 | 2.9 | 0 | 11 | 0.33 |
| *ESS* | 5.52 | 3.82 | 0 | 13 | 7.18 | 3.94 | 0 | 14 | 0.15 |
| *PSQI* | 3.95 | 2.22 | 0 | 8 | 4.65 | 1.95 | 1 | 9 | 0.24 |
| *Sex (F - M)* | 14 F – 5 M |  |  |  | 29 F – 5 M |  |  |  | 0.15 |

The education level is expressed in years of study. BMI stands for Body Mass Index and is expressed in kg/m<sup>2</sup>. BDI score, BAI score, ESS score and PSQI respectively stand for Beck Depression Inventory score (Beck, Steer and Carbin 1988), Beck Anxiety Inventory score (Beck, Epstein, et al. 1988), Epworth Sleepiness Scale score (Johns 1993) and Pittsburgh Sleep Quality Index (Buysse, et al. 1989). F: Female and M: Male. All characteristics were compared using two-sample *t*-tests, except for sex composition that were compared using a Chi-square test.

**Table 2:** Activation foci within the LC at the appearance of target sounds, for a statistical threshold of p < 0.05 FWE-corrected after small-volume correction using the LC template as search region. R: Right, L: Left.

| Brain region<br>(Peak MNI coordinates) | Cluster size | T<sub>49</sub> | p-FWE | MNI coordinates<br>(x, y, z) in mm |  |  |
| --- | --- | --- | --- | --- | --- | --- |
| Locus coeruleus L | 25 | 4.93 | 0.001 | -4 | -36 | -20 |
|  | 6 | 4.87 | 0.001 | -8 | -38 | -32 |
| Locus coeruleus R | 2 | 4.42 | 0.003 | 8 | -37 | -32 |
|  | 4 | 3.57 | 0.03 | 6 | -35 | -28 |
|  | 2 | 3.4 | 0.046 | 4 | -36 | -19 |

The exclusion criteria were as follows: history of major neurologic or psychiatric disease or stroke; recent history of depression and anxiety (< 5 years); sleep disorder; use of any medication affecting the central nervous system; smoking; excessive alcohol (> 14 units/week) or caffeine (> 5 cups/day) consumption; night shift work in the past 6 months; travels in a different time zone during the last 2 months; Body Mass Index (BMI) ≤ 18 and ≥ 29 (for the older participants) and ≥ 25 (for the younger participants); clinical symptoms of cognitive impairment for older subjects (Mattis Dementia Rating Scale score < 130; Mini-Mental State Examination score < 27) (Coblentz et al., 1973; Folstein et al., 1975) and MRI contraindications. Due to miscalculation at screening, one older participant had a BMI of 30.9 and one of the younger participants had a BMI of 28.4. Since their BMI did not deviate substantially from the criteria and that BMI was used as a covariate in our statistical models, these participants were included in the analyses. Depression, anxiety, sleepiness, and sleep quality were assessed with the Beck Depression Inventory (BDI) (Beck, Steer, et al., 1988), Beck Anxiety Inventory (BAI) (Beck, Brown, et al., 1988), Epworth Sleepiness Scale (ESS) (Johns, 1993) and the Pittsburgh Sleep Quality Index (PSQI) (Buysse et al., 1988), respectively.

The younger participants were requested to maintain a loose fixed sleep-wake schedule (+- 1h) for 1 week before fMRI acquisitions to reduce prior sleep deprivation and favor similar circadian entrainment across participants while keeping realistic daily life conditions. Adherence to the schedule was verified using a wrist actimetry device (AX3, Axivity Ltd, Newcastle, UK). Older participants were requested to avoid unusual late sleep time for 3 days prior to their participation. Adherence was verified using sleep diaries.

FMRI recordings were completed in the morning, 2 to 3 h after wake-up time to control for time-of-day effects. Prior to entering the MRI scanner, all participants were maintained in dim light (10 lux) for at least 45 min during which they received instructions about the following MRI sessions. The fMRI sessions consisted of a 10 min visual task followed by a 10 min auditory oddball task. The present paper only deals with the auditory task. Structural MRI data were acquired in a separate MRI session completed within 1 week before or after the fMRI session.

### Auditory oddball task

The task consisted of rare deviant target tones (1000 Hz sinusoidal waves, 100 ms) composing 20% of the tones that were pseudorandomly interleaved within a stream of standard stimuli (500 Hz sinusoidal waves, 100 ms). The task included 270 auditory stimuli in total (54 target tones). Auditory stimuli were delivered to the participants with MRI-compatible headphones (Sensimetrics, Malden, MA). The interstimulus interval was set to 2000 ms. Participants were instructed to press with the right index finger on an MRI-compatible keyboard (Current Designs, Philadelphia, PA) as quickly as possible at the appearance of target sounds. The experimental paradigm was designed using OpenSesame 3.2.8 (Mathôt et al., 2012). The MRI session started with a short volume calibration session to ensure an optimal perception of the stimuli.

### MRI data acquisitions

MRI data were acquired using a MAGNETOM Terra 7T MRI system (Siemens Healthineers, Erlangen, Germany), with a single-channel transmit and 32 channel receive head coil (1TX/32RX Head Coil, Nova Medical, Wilmington, MA). To reduce dielectric artifacts and homogenize the magnetic field of Radio Frequency (RF) pulses, dielectric pads were placed between the head of the participants and the coil (Multiwave Imaging, Marseille, France).

BOLD fMRI data were acquired during the task, using the CMRR multi-band (MB) Gradient-Recalled Echo - Echo-Planar Imaging (GRE-EPI) sequence: TR = 2340 ms, TE = 24 ms, flip angle = 90°, matrix size = 160 × 160, 86 axial 1.4mm-thick slices, MB acceleration factor = 2, GRAPPA acceleration factor = 3, voxel size = (1.4×1.4×1.4)mm^3^. The cardiac pulse and the respiratory movement were recorded concomitantly using a pulse oximeter and a breathing belt (Siemens Healthineers, Erlangen, Germany), respectively. The fMRI acquisition was followed by a 2D GRE field mapping sequence to assess B0 inhomogeneity with the following parameters: TR = 5.2 ms, TEs = 2.26 ms and 3.28 ms, FA = 15°, bandwidth = 737 Hz/pixel, matrix size = 96 × 128, 96 axial 2.0mm-thick slices, voxel size = (2×2×2)mm, acquisition time = 1:38 min.

A Magnetization-Prepared with 2 RApid Gradient Echoes (MP2RAGE) sequence was used to acquire T1-weighted anatomical images: TR = 4300 ms, TE = 1.98 ms, FA1/FA2 = 5°/6°, TI1/TI2 = 940 ms/2830 ms, bandwidth = 240 Hz/pixel, matrix size = 296 × 256, 224 axial 0.75mm-thick slices, GRAPPA acceleration factor = 3, voxel size = (0.75×0.75×0.75)mm^3^, acquisition time = 9:03 min (Marques & Gruetter, 2013). The LC-specific sequence consisted of a 3D high-resolution Magnetization Transfer-weighted Turbo-FLash (MT-TFL) sequence with the following parameters: TR = 400 ms, TE = 2.55 ms, FA = 8°, bandwidth = 300 Hz/pixel, matrix size = 480 × 480, number of averages = 2, turbo factor = 54, MTC pulses = 20, MTC FA = 260°, MTC RF duration = 10000 μs, MTC Inter RF delay = 4000 μs, MTC offset = 2000 Hz, voxel size = (0.4×0.4×0.5)mm^3^, acquisition time = 8:13 min. Sixty axial slices were acquired and centered for the acquisitions perpendicularly to the rhomboid fossa (i.e., the floor of the fourth ventricle located on the dorsal surface of the pons).

### MRI data pre-processing

EPI images were realigned and unwarped using the Statistical Parametric Mapping toolbox (SPM12, https://www.fil.ion.ucl.ac.uk/spm/software/download/). EPI images underwent brain extraction using “BET” from the FMRIB Software Library suite (FSL, https://fsl.fmrib.ox.ac.uk/fsl/fslwiki) and the final images were spatially smoothed with a Gaussian kernel characterized by a full width at half maximum of 3 mm.

The background noise in MP2RAGE images was removed using an extension of SPM12 (extension : https://github.com/benoitberanger/mp2rage) (O’Brien et al., 2014). The denoised image was then automatically reoriented using the *‘spm-auto-reorient’* SPM function and corrected for intensity non-uniformity using the bias correction method implemented in the SPM segmentation. Brain extraction was then conducted on the denoised-reoriented-biased-corrected image using both the Advanced Normalization Tools (ANTs, http://stnava.github.io/ANTs/) (Avants et al., 2011) with the *‘antsBrainExtraction’* function and the RObust Brain EXtraction tool (ROBEX, https://www.nitrc.org/projects/robex) (Iglesias et al., 2011). The method yielding to the best extraction for each individual as assessed by visual inspection, was used for subsequent steps. A whole-brain T1 group template was created using ANTs, based on preprocessed MP2RAGE images of all subjects except for one, the MP2RAGE image of whom was not adapted due to a bad positioning of the slices during the acquisitions. Finally, the preprocessed MP2RAGE image of each subject was normalized to the Montreal Neurological Institute (MNI) space (with a 1×1×1 mm^3^ image resolution). The purpose of using a template that is specific to our dataset, was to improve the registration into the MNI space using an intermediate space. The transformation parameters obtained from normalization were later used for registering first-level statistical maps into the MNI space to conduct group-level fMRI analyses (see Statistical Analyses section).

In order to extract LC contrast, T1 structural images in the space of the subject (after removing the background noise) were upsampled by a factor 2 [(0.375×0.375×0.375)mm^3^] to avoid losing in-plane resolution when registering the LC slab to the T1 image. The upsampling was done using the ‘*nii_scale_dims’* function from an extension of SPM12 (extension : https://github.com/rordenlab/spmScripts). The complete LC contrast extraction was done in the original space of the subject. The MT-TFL image of each subject was registered with the whole brain upsampled T1 image by means of a two-step process: (i) a rough manual registration to extract the parameters for an initial transformation using ITK-SNAP (Yushkevich et al., 2016), and (ii) an automatic affine registration based on the initial transformation parameters, using ANTs. MT-TFL data of one subject was not usable, due to an excessive motion of the participant, leading to a registration failure. The LC appearing hyperintense on registered MT-TFL images was manually delineated by two expert raters and the intersection of the LC masks of the two raters was computed as the final LC mask for each individual. The LC mask was skeletonized by only keeping the voxel with the highest intensity in each axial slice. Based on the skeletonized LC mask, the LC contrast was computed after normalization of each LC slice intensity to a slice-corresponding 2D reference region (a 15 × 15 voxels region, corresponding to a (5.5 × 5.5)mm^2^ square region) situated anteriorly (and centrally) in the pons, in the pontine tegmentum. For example, the left LC contrast was defined as:

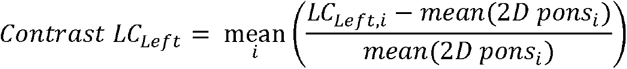

Where:

- i is the slice index along the (left) LC

*- LC*_*Left,i*_ is the intensity of the voxel with the highest intensity in the axial slice with index i

- mean(2D pons_i_) represents the mean intensity in the 2D reference region corresponding to the axial slice with index i

The LC contrast was computed as the mean LC contrast between the left and right LC. The distribution of the LC contrasts across individuals was investigated by computing the Probability Density Function (PDF), using a kernel density nonparametric method (*ksdensity* MATLAB R2021a built-in function). Individual skeletonized LC masks were used for extracting the LC activity during the oddball task in the subject space. In order to investigate the activation of the LC at the group-level, an LC probabilistic template was created. The LC mask of each volunteer was normalized to the structural group template, and then to the MNI (MNI152 - 1×1×1mm^3^). This was done using the *‘antsApplyTransforms’* ANTs command, with the transformation parameters estimated (i) when registering the subject-specific MP2RAGE image to the structural template, and (ii) the transformation parameters estimated when registering the structural template to the MNI. The final LC probabilistic template was created as the sum of all masks divided by the number of subjects included in the analysis.

### Statistical Analyses

Statistical analyses were conducted using SPM12. A high-pass filter with a 128 s cutoff was applied to remove slow signal drifts. The timing vector with the appearance of the target tones was convolved with the canonical Hemodynamic Response Function (HRF) to model the event-related response and was used as the main condition in a General Linear Model (GLM). The PhysIO Toolbox (https://www.tnu.ethz.ch/en/software/tapas/documentations/physio-toolbox) was used to compute physiology-related voxel-wise signal fluctuations based on respiratory and cardiac pulsation data (Kasper et al., 2017), that was available in 48 volunteers (physiological data was not available for 5 volunteers). The Fourier expansion of cardiac and respiratory phase computed with the toolbox as well as the realignment parameters were used as multiple regressors of no interest in the GLM. To avoid any registration-induced error, the first-level statistical analysis was conducted in the space of the subject.

The mean functional image was registered to the MP2RAGE image to extract the corresponding transformation matrix used to register the first-level statistical map of each subject to the structural image. Therefore, for all subjects, statistical maps corresponding to the appearance of target sounds were registered to the native space, normalized to the group template space and then to the MNI space. A second-level analysis was then conducted in the MNI space (to report coordinates of activation clusters), where age, sex and BMI were used as covariates. Whole brain activation was first assessed following voxel-level Family-Wise Error (FWE) correction based on random field theory for p < 0.05. For the sake of simplicity, only clusters with a size of at least 20 voxels were reported (an extensive table without minimum cluster size and all significant voxel cluster is available upon request to the corresponding author). Structures showing peak activation foci were identified using the Harvard-Oxford Subcortical and Cortical structural Atlases (Desikan et al., 2006). The probabilistic mask of the LC was then used to assess specific activation of the LC. Due to the small size of the nucleus, LC activation was not expected to survive stringent whole brain FWE correction. Therefore, a small-volume correction using the LC template was conducted using SPM12 to report voxel-level FWE-corrected results within the LC mask.

REX Toolbox (https://web.mit.edu/swg/software.htm) was then used to extract the activity estimates (betas) associated with the appearance of the target sounds in the skeletonized LC mask of each subject, within the subject space (Duff et al., 2007). This procedure ensured that any potential displacement and bias introduced by the normalization step into the common MNI space did not affect individual activity estimates. Statistical analyses using these activity estimates were performed in Rstudio (version 2022.07.1; https://www.rstudio.com/). For all models, a multivariate linear modeling approach was used, using sex, BMI and education as covariates. LC response and contrast followed Gaussian distributions. The three models of interest investigated in the present study were :

i. LC_contrast_∼ age + covariates, (ii) LC_response_∼ age + covariates and (iii) LC_response_∼ LC_contrast_ + age + (LC_contrast_ x age) + covariates. The first and second models were designed to assess age-related changes in LC contrast and LC response, respectively. The third model was intended to seek a relationship between functional response and LC contrast.

## RESULTS

The LC contrast across the entire length of the nucleus was extracted for each individual based on MT-TFL images. The multivariate linear model using the LC contrast averaged over both LCs as dependent variable found a significant main effect of age (**p = 0.038**), but no main effect for BMI (p = 0.456), sex (p = 0.565) and education (p = 0.943) (**Figure 1A**). When adding age-group in addition to age to the model to take into account the fact that age was not truly continuous in our sample, these statistical outputs remain significant (main effect of age: p = 0.012; main effect of age-group: p = 0.035). The LC contrast distributions in young and late middle-aged individuals are shown in **Figure 1B**. The same model was then computed for the left LC contrast (p = 0.053) and the right LC contrast (p = 0.15), separately. We further explored potential differences in LC contrast in late middle-aged individuals aged 60 or less and 61 or more in a multivariate linear model similar to the preceding and found no main effect of age subgroups (p = 0.16). The LC contrast distributions in late middle-aged subgroups are shown in **Figure 1C**.

**Figure 1:**
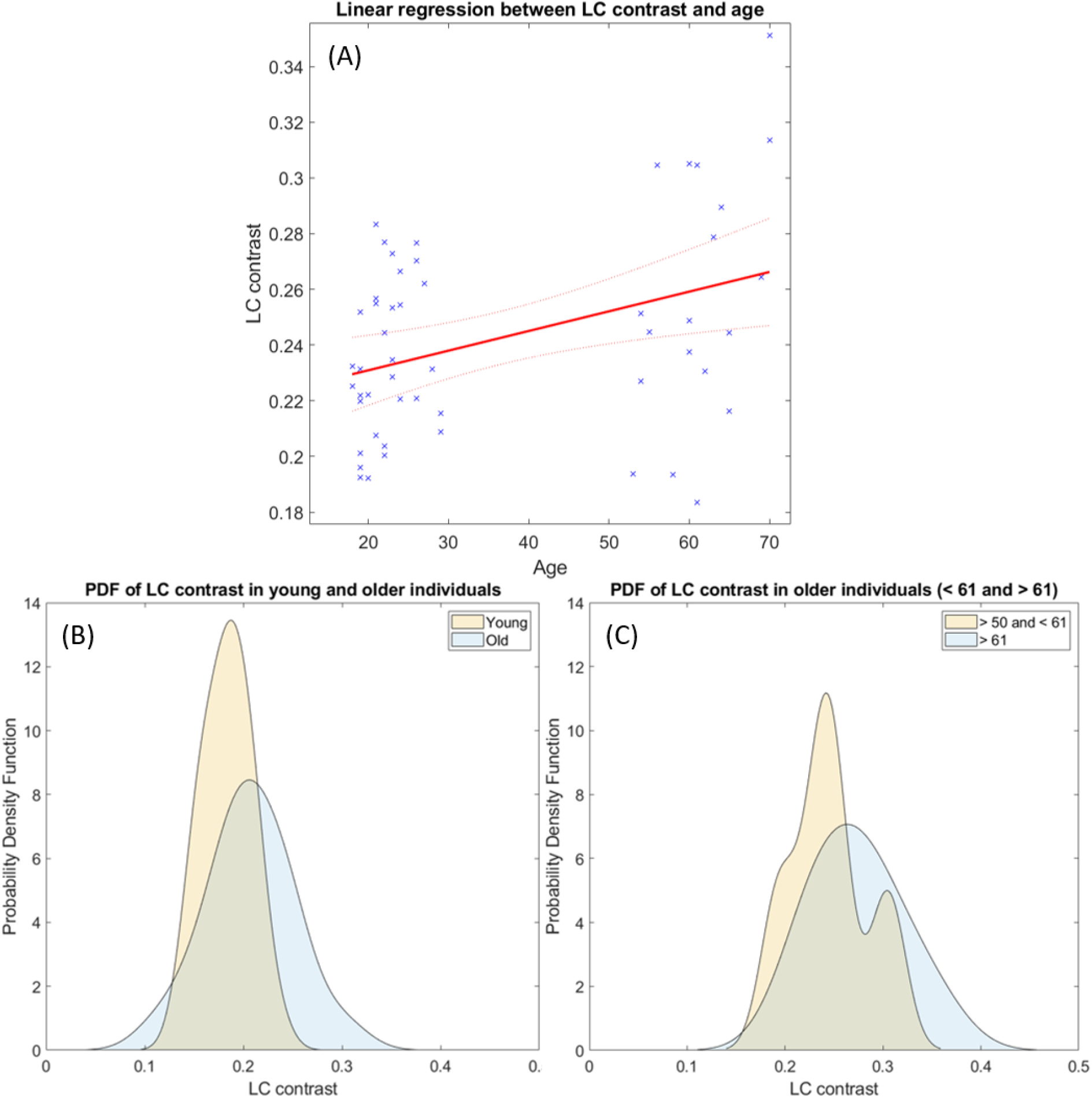
LC contrast variation with age. (A) Linear regression plot of LC contrast and age. Solid line: regression line; Dashed lines: 95% confidence interval (B) Probability Density Function (PDF) of the LC contrast for young and older participants, (C) and subgroups of the older cohort (< 61 and > 61 y).

During the fMRI recordings, the participants completed the task correctly (mean accuracy: 97.1%, SD: 10.4%). FMRI data analyses over the entire brain showed that the target tones were associated with a significant greater BOLD signal in a wide set of areas (**Figure 2**). At the cortical level, significant activation foci (p_FWE_ < 0.05) were detected bilaterally over the cerebellum, the posterior cingulate gyrus, the insular cortex, the precuneus, the middle temporal gyrus, the middle frontal gyrus, the frontal pole, and unilaterally over the anterior cingulate gyrus (left), the planum polare (left), the lateral occipital cortex (right), the cuneal cortex (left), the superior frontal gyrus (left), the precentral gyrus (left) and the lingual gyrus (right) (**Figure 2A**). At the subcortical level, a significant activation was detected bilaterally over the thalamus and the caudate. These results are in line with the reported neural correlates of mismatch negativity tasks and support the validity of our procedure (Brázdil et al., 2005; Kiehl et al., 2001a, 2001b; Linden et al., 1999; Menon et al., 1997). A detailed list of brain activations highlighted using a stringent statistical threshold controlling for multiple comparison is provided in the supplementary materials and may serve as a reference for future 7T MRI investigations using the same task (**Supplementary Table S1**). No significant activation difference was observed between healthy young and late-middle-aged individuals (for a statistical threshold p_FWE_ < 0.05).

**Figure 2:**
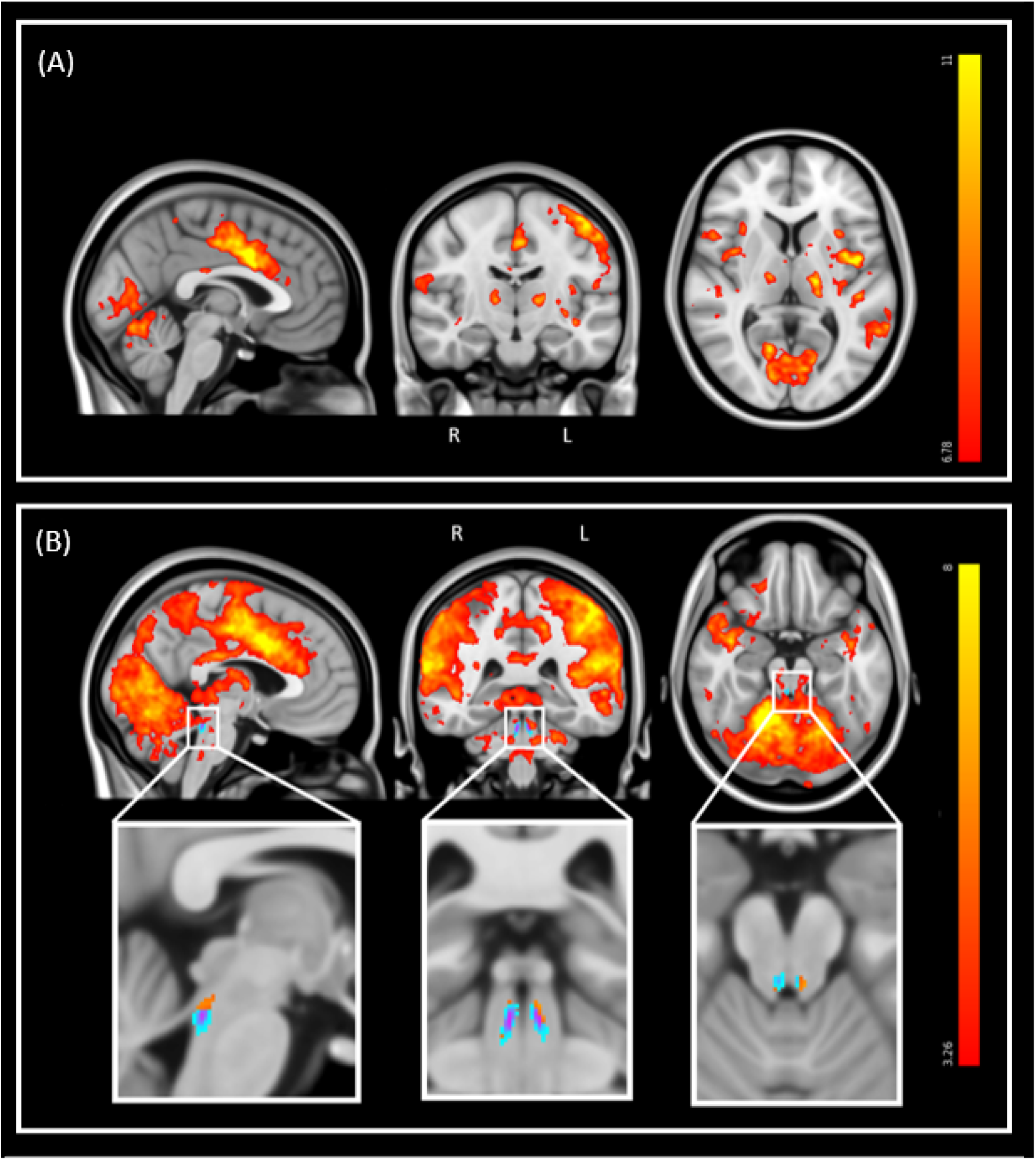
Whole-brain and LC response to the target sounds during the auditory oddball task. Sagittal, coronal, and axial views (MNI coordinates: [-3 -15 7]). The legend shows the t-values associated with the color maps. The images are shown with the radiological convention for the orientation. (A) Whole-brain results using significance for a threshold of p < 0.05 FWE-corrected (p < 3.34e-8 uncorrected ; t > 6.35), and a minimum cluster size of 20 voxels. Refer to table S1 for detailed list of coordinates. (B) Same results displayed using an uncorrected threshold of p < 0.001 (t > 3.26). Insets at the bottom show the LC probabilistic template and the significant activation detected within this mask (p < 0.05 FWE-corrected within LC mask). Refer to table 2 for detailed coordinates.

We then focused on the LC. Given the small size of the structure we did not expect that an activation would survive the conservative p_FWE_ < 0.05 statistical threshold over the whole brain. Correction for multiple comparisons p_FWE_ < 0.05 was rather considered within the study-specific probabilistic template of the LC in the MNI space. Following this procedure, five significant local peak intensity clusters were detected in the rostral part of the LC (**Table 2**). Although a significant activation was detected bilaterally, the activation clusters were larger over the rostral left LC (**Figure 2B**).

Further analyses were conducted in the native space of each subject, where the individual LC activity was extracted. The multivariate linear model for the bilaterally averaged estimates of the LC activity did not yield significant main effects for age (p = 0.57), BMI (p = 0.13), sex (p = 0.49) or education (p = 0.7). Similar statistical outputs were obtained when computing the same model separately with the activity estimate of the left and right LC (main effects for age: left LC, p = 0.92, right LC, p = 0.3) or when adding age group to the model (main effect for age and group: left LC, p > 0.3, right LC, p > 0.5). We then used a multivariate linear model to assess the link between LC activity, as dependent variable, and LC contrast. The model did not reveal a significant main effect of LC contrast (p = 0.77), nor an LC contrast by age interaction (p = 0.77) on the LC response (**Figure 3**). Similar statistical outputs were obtained when computing the same model separately with the activity estimate and contrast of the left and right LC, respectively (main effects for LC contrast - left LC : p = 0.63; right LC : p = 0.16), or when adding age group to the model (main effect for age and group: left LC, p > 0.4, right LC, p > 0.3)

**Figure 3:**
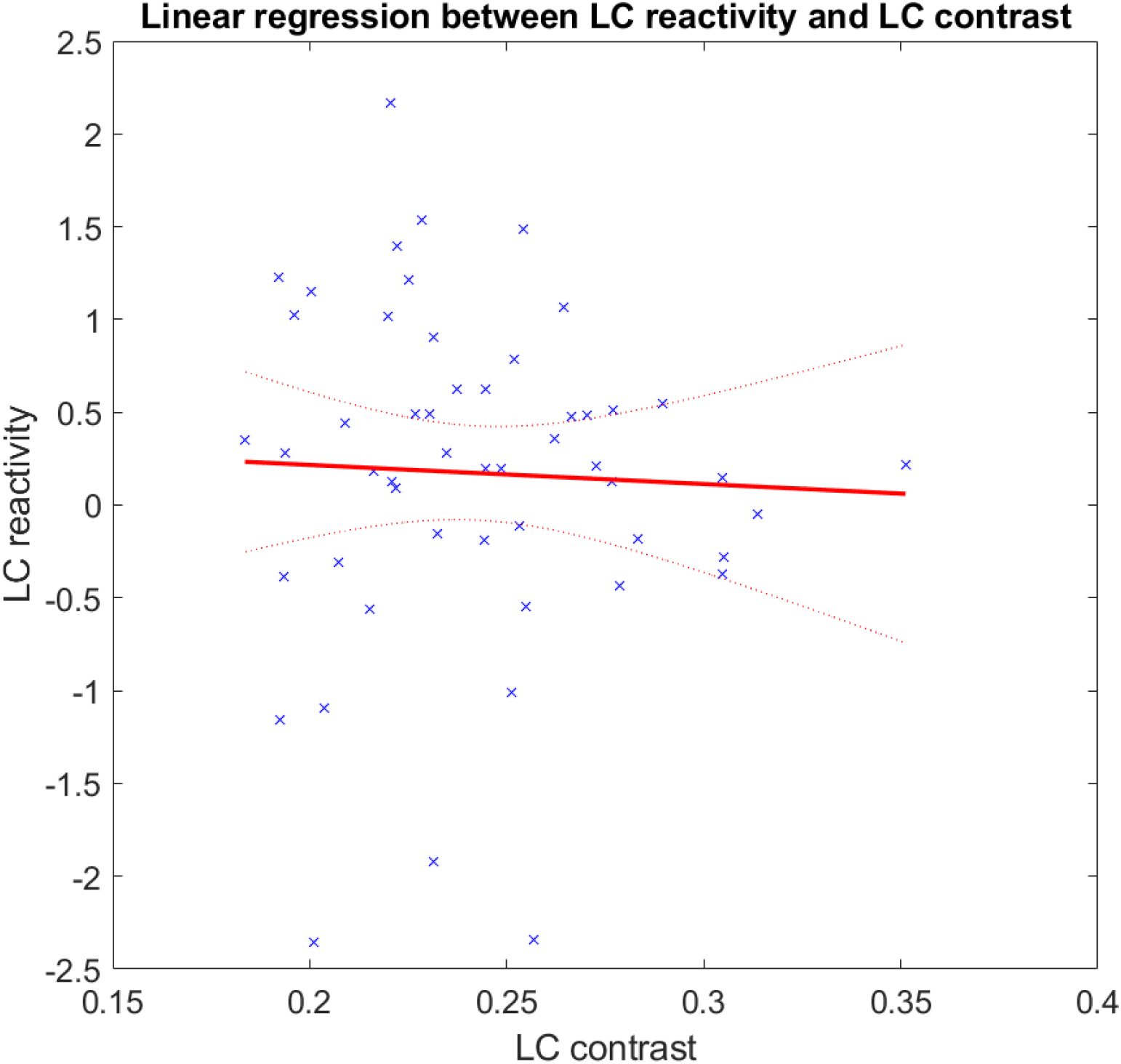
Association between LC activation and LC contrast. Linear regression plot of LC activity and LC contrast. Solid line: regression line; Dashed lines: 95% confidence interval.

## DISCUSSION

A better characterization of structural and functional characteristics of the LC in healthy individuals is critical (i) to detect abnormal early features occurring in an array of diseases and disorders affecting the LC, (ii) to characterize their possible evolution, and (iii) potentially help in the development of therapies targeting the LC-NE system. We assessed the LC functional response and LC contrast in a cohort of 53 healthy individuals aged 18 to 30 years and 50 to 70 years using a UHF 7T MRI system. We did not find an association between age and LC response, as assessed during an auditory oddball task. While we found the expected age-related increase in LC contrast, this accepted marker of LC integrity was not significantly related to its functional response.

Although widely studied, the exact source of the LC contrast as extracted using specific MRI sequences remains debated (Galgani et al., 2021). It was hypothesized that LC contrast was related to the progressive increase in neuromelanin that accumulates inside the cell bodies of noradrenergic neurons, as confirmed in *post-mortem* studies (Fedorow et al., 2006; Zucca et al., 2006). Histology combined with *post-mortem* 7T MRI acquisitions, revealed that regions with T1-hyperintensities within the LC colocalized with the presence of neuromelanin-rich neurons (Keren et al., 2015) such that LC contrast might reflect the density of neuromelanin-containing noradrenergic neurons within the LC (Betts et al., 2017; Liu et al., 2019). Recent investigations further showed that the LC contrast obtained using a T1-weighted MRI sequence with magnetization transfer was still detectable in mice genetically modified to have LC damages (leading to LC cell loss) (Watanabe et al., 2019). Therefore, the LC contrast may not be directly linked to the accumulation of neuromelanin itself, but rather to the specific microstructure of noradrenergic neurons (Watanabe et al., 2019). Overall, while the LC contrast is deemed to reflect the integrity of the LC (i.e. its neuronal density), lipid accumulation and inflammation have also been suspected to influence the LC contrast (Priovoulos et al., 2020).

In line with the literature, an expected age-related increase in LC contrast was observed in the present study (Betts et al., 2017; Clewett et al., 2016). A previous study hypothesized that the increased LC contrast with age may be linked to a related shrinkage of cells within the LC (Liu et al., 2019). However, a study using staining methods also indicated that the volume of the LC and its neuron population were not affected by normal aging (age range: 47-83 years) (Theofilas et al., 2017). We found no indication of a potential plateau or a decreased LC contrast starting after 60 years in our sample, that would be compatible with the previously reported inverted-U relationship between contrast and age with a maximum contrast found at around 60 years (Liu et al., 2019; Shibata et al., 2006). This may be due to a lack of sensitivity as our sample of late middle-aged individuals was relatively young and small : 19 individuals in total with only 10 participants with > 60 years, and a range of ages 50-70 years. This may also come in part from the stringent exclusion criteria which led to a sample of very healthy individuals in which LC integrity could presumably be better preserved. This does not mean, however, that they could not harbor presymptomatic pathologies such as tauopathy (Liu et al., 2019). For example, a lower LC contrast is related to higher tau deposition in the entorhinal cortex in cognitively unimpaired individuals (Jacobs et al., 2021). Therefore, when studying the effect of normal aging, one cannot exclude that participants included are in a prodromal phase of age-related diseases. Indeed, based on autopsy data, knowing that approximately 50% of individuals between 30 and 40 years present an accumulation of abnormal tau proteins within the LC (Braak et al., 2011; Braak & del Tredici, 2011), some participants in the current study could already be in a Braak stage I/II. However, a tau PET examination would be needed to determine the Braak stage in our cohort of cognitively intact participants.

The LC is known to be involved in novelty or salience detection (Krebs et al., 2018; Murphy et al., 2014; Sterpenich et al., 2006). It is reciprocally connected with the cortical salience network to enhance the processing of behaviorally important stimuli (Lee et al., 2020). The salience network is involved in high-level cognitive control to detect unexpected stimuli and reorient attention (Bouret & Sara, 2005). In the present study, a reliable LC response was detected with UHF 7T MRI system during an auditory oddball paradigm, which mimics novelty and/or salience detection. In contrast to a prior study (Murphy et al., 2014) where pupil response – an accepted output of LC phasic activity – was recorded and used as a regressor to isolate LC response during an auditory oddball task, no pupil measurement was required in the present study to extract the LC response. We detected a bilateral activation in the rostral LC – a part of the nucleus densely connected to associative regions (Jacobs et al., 2020) – that was mostly left lateralized. While there is no clear consensus on the presence of a lateralization in the LC, there is increasing evidence of regional specialization of the LC in terms of cell composition and projections (Poe et al., 2020). Previous tracing studies conducted in rats showed that rostral LC projections innervate a multitude of brain areas, including the hippocampus, the septum, the caudate-putamen (Mason & Fibiger, 1979) and the hypothalamus (Loughlin et al., 1986; Mason & Fibiger, 1979). While an intense labeling was observed in the caudal LC after injection of tracers into the thalamus, a scattered cell labeling in the rostral LC was also reported (Mason & Fibiger, 1979; Simpson et al., 1997). Although translation to humans may not be straightforward, these results support that the LC subparts are not uniform in terms of projections to the different brain areas. It is therefore plausible that the rostral activation reflects, at least in part, a true functional regional effect related to the ongoing cognitive process during the oddball task, rather than to sensitivity issues (that we cannot, however, rule out).

The fact that the LC response remained stable in late middle-aged individuals compared to young adults was not in line with our expectations. Once again, these results could be due to the very healthy and relatively young nature of our sample or the inability of fMRI to capture subtle LC response differences between individuals during an auditory oddball task. The oddball task may also be too cognitively undemanding to trigger true age-related differences. Despite this, our results suggest that the age-related increase in LC contrast does not necessarily translate in a detectable change in functional response. This observation is further reinforced by the absence of correlation between LC contrast and LC response - at least in a simple auditory mismatch negativity task. Although longitudinal data could confirm this causality, our results may support the idea that LC contrast is more sensitive to aging or takes place before functional changes. One could hypothesize that a compensation mechanism takes place to counter age-related alterations in the LC structure, as reflected by a change in LC contrast. This compensation mechanism would, however, need to scale out exactly the consequence of the structural alteration so that no difference between age groups can be detected. This cannot be ruled out though compensatory mechanisms to sustain cognition in aging most often result in higher activation or bilateralization of brain responses (Reuter-Lorenz & Park, 2010).

The activity of the LC neurons is known to follow tonic and phasic modes. While the tonic activity of LC neurons is known to be involved in task disengagement and search of alternative behaviors (exploration), the phasic activity during a goal-oriented task facilitates task-related behaviors in order to maximize the performance (exploitation) (Aston-Jones & Cohen, 2005). A trade-off between these two modes allows to maximize the reward and the utility (Aston-Jones & Cohen, 2005). Since the auditory oddball paradigm in the present study did not require the search for alternative behaviors at the expense of a task performance optimization, one could hypothesize that only the phasic activity of the LC was investigated with our protocol. The repetition time used for our acquisitions (i.e. 2.34 s) is, however, relatively long compared to the fast burstiness of LC neurons, such that tonic activity or interactions between tonic and phasic activity could contribute to our findings. However, future studies should evaluate the relationship between age-related changes in LC contrast and tonic activity. Other analyses could also probe a potential link between LC contrast and resting-state functional connectivity using the LC as a seed region (Jacobs et al., 2018).

It is worth mentioning that while linear patterns are sought in the present study, no individual between the age of 30-50 was sampled, constituting a limitation. A study with a less discretized data sample could confirm our results. Future studies may also want to use individually tailored Hemodynamic Response Functions (HRF) to assess LC response. While the canonical HRF we used to model activity over entire brain seems suitable to model average LC response over a group of participants, individual LC responses can vary substantially across individuals (Prokopiou et al., 2022b).

Given the critical involvement of the LC in cognitive and behavioral processes such as learning and memory (Kety, 1972; Sara, 1985), attention (Sara & Segal, 1991), regulation of sleep and vigilance (Aston-Jones & Bloom, 1981), or addiction (Aghajanian, 1978), assessing its response could be useful to evaluate the integrity of the LC-NE system in a vast array of psychiatric, neurologic, and neurodegenerative disorders. For example, alteration in the activity and/or connectivity of the LC were reported to contribute to the development of depression (del Cerro et al., 2020; Szot et al., 2016) and the symptomatology of schizophrenia (Yamamoto et al., 2014). Aside from the early tauopathy of the LC that appears to contribute to AD (Jacobs et al., 2021), synucleinopathy contributing to LC degeneration was also detected in prodromal dementia of Lewis bodies (Hansen, 2021). Finally, concerning existing therapies targeting the LC-NE system, it was suggested that the functional integrity of this system may be linked to responsiveness to Vagus Nerve Stimulation in patients with inoperable drug-resistant epilepsy (Berger et al., 2021; de Taeye et al., 2014; Hödl et al., 2020). In the context of healthy aging, our results indicate that, while LC contrast changes with age, LC responsiveness may remain stable such that any abnormal response may constitute an early sign of an unhealthy trajectory.

## Supporting information

Supplementary_matrials

## Acknowledgments

We thank E. Lambot, C. Hagelstein, B Lauricella, P. Hawotte, A. Claes, B. Herbillon, P. Maquet, C. Bastin, F. Collette, E. Salmon, M. Bahri, N. Belly, G. Hammad for their help in the different steps of the project.

