## Supplementary_matrials for "MRI-assessed locus coeruleus contrast and functional response are not associated in young and late middle-aged individuals"

***Supplementary materials***

***Table S1: Significant responses to the target sound.***

| **Brain region**  **(Peak MNI coordinates)** | **Cluster size** | **T_49_** | **MNI coordinates**  **(x, y, z) in mm** | | |
| --- | --- | --- | --- | --- | --- |
| Insular Cortex L | 59040 | 15.4465 | -41 | -4 | 11 |
| Cerebellum (VI) R | 34354 | 17.781 | 30 | -55 | -26 |
| Middle Temporal Gyrus (TOP) R | 18333 | 12.5287 | 63 | -48 | 10 |
| Cingulate Gyrus (AD) L | 16879 | 17.86 | -1 | 5 | 36 |
| Insular Cortex R | 12563 | 12.1277 | 42 | 5 | -2 |
| Thalamus L | 1549 | 13.3082 | -16 | -18 | 5 |
| Middle Temporal Gyrus (PD) R | 972 | 10.6742 | 48 | -25 | -6 |
| Cingulate Gyrus (PD) R | 754 | 9.6633 | 6 | -19 | 26 |
| Thalamus R | 541 | 10.0317 | 15 | -14 | 7 |
| Frontal Pole R | 466 | 7.4437 | 39 | 41 | 22 |
| Precuneus Cortex L | 194 | 8.6936 | -1 | -44 | 59 |
| Thalamus R | 187 | 9.0946 | 15 | -6 | 9 |
| Middle Frontal Gyrus R | 145 | 8.2982 | 38 | -1 | 58 |
| Middle Temporal Gyrus (PD) L | 132 | 8.7711 | -66 | -29 | -3 |
| Precuneus Cortex L | 96 | 7.8525 | -9 | -49 | 50 |
| Planum Polare L | 95 | 8.621 | -40 | -2 | -18 |
| Middle Temporal Gyrus (TOP) R | 93 | 8.1126 | 46 | -37 | 0 |
| Middle Frontal Gyrus L | 92 | 7.6677 | -31 | 37 | 39 |
| Frontal Pole L | 91 | 7.4702 | -28 | 54 | 22 |
| Frontal Pole R | 82 | 9.5149 | 25 | 41 | -17 |
| Cerebellum (VI) L | 70 | 7.6427 | -8 | -74 | -24 |
| Caudate R | 62 | 7.8333 | 14 | 15 | 4 |
| Middle Temporal Gyrus (PD) L | 58 | 7.5001 | -50 | -30 | -6 |
| Thalamus R | 50 | 8.7706 | 4 | -5 | 7 |
| Frontal Orbital Cortex R | 48 | 8.044 | 29 | 19 | -13 |
| Middle Temporal Gyrus (PD) L | 47 | 7.1467 | -59 | -28 | -4 |
| Precuneus Cortex R | 46 | 7.0337 | 7 | -50 | 47 |
| Lateral Occipital Cortex (SD) R | 42 | 7.0278 | 35 | -61 | 42 |
| Cingulate Gyrus (PD) R | 40 | 6.9469 | 13 | -23 | 36 |
| Caudate R | 39 | 7.4336 | 12 | 6 | 9 |
| Cerebellum (VIIIa) R | 37 | 8.6327 | 9 | -67 | -49 |
| Precuneus Cortex R | 34 | 7.8746 | 8 | -46 | 53 |
| Superior Frontal Gyrus L | 31 | 7.7228 | -25 | 4 | 65 |
| Cerebellum (I-IV) L | 27 | 7.2685 | -7 | -46 | -19 |
| Frontal Pole L | 27 | 6.6931 | -35 | 50 | 22 |
| Cuneal Cortex L | 25 | 7.1918 | 0 | -89 | 15 |
| Precentral Gyrus L | 23 | 7.2071 | -52 | 1 | 34 |
| Lingual Gyrus R | 22 | 7.1215 | 14 | -68 | -5 |

*Significant activation foci for a statistical threshold of p < 0.05 FWE-corrected (p < 3.34e-8 uncorrected; t > 6.35), and a minimum cluster size of 20 voxels (38 clusters reported). R: Right, L: Left, TOP: Temporo-Occipital Part, AD: Anterior Division, PD: Posterior Division, SD: Superior Division. An extensive table without minimum cluster size and including 138 significant voxel clusters is available upon request to the corresponding author.*
